# DR2S: An Integrated Algorithm Providing Reference-Grade Haplotype Sequences from Heterozygous Samples

**DOI:** 10.1101/2020.11.09.374140

**Authors:** Steffen Klasberg, Alexander H. Schmidt, Vinzenz Lange, Gerhard Schöfl

## Abstract

**Background:** High resolution *HLA* genotyping of donors and recipients is a crucially important prerequisite for haematopoetic stem-cell transplantation and relies heavily on the quality and completeness of immunogenetic reference sequence databases of allelic variation.

**Results:** Here, we report on `DR2S`, an R package that leverages the strengths of two sequencing technologies – the accuracy of next-generation sequencing with the read length of third-generation sequencing technologies like PacBio’s SMRT sequencing or ONT’s nanopore sequencing – to reconstruct fully-phased high-quality full-length haplotype sequences. Although optimised for *HLA* and KIR genes, `DR2S` is applicable to all loci with known reference sequences provided that full-length sequencing data is available for analysis. In addition, `DR2S` integrates supporting tools for easy visualisation and quality control of the reconstructed haplotype to ensure suitability for submission to public allele databases.

**Conclusions:** `DR2S` is a largely automated workflow designed to create high-quality fully-phased reference allele sequences for highly polymorphic gene regions such as *HLA* or *KIR*. It has been used by biologists to successfully characterise and submit more than 500 *HLA* alleles and more than 500 *KIR* alleles to the *IPD-IMGT/HLA* and *IPD-KIR* databases.

## Background

The *human leukocyte antigen* (*HLA*) genes encode key constituents of the human adaptive immune system and are amongst the most polymorphic genes of the human genome [1]. Currently (Jan. 2021), the public immunopolymorphism database *IPD-IMGT/HLA* lists 27,059 different alleles for the six ”classical” *HLA* genes (*A, B, C, DRB1, DQB1, DPB1*) alone, and it is still growing substantially at each quarterly release [2]. For haematopoetic stem-cell transplantation (*HSCT*), allelic matching between patients and donors for these *HLA* genes is a key determinant of success [3], as each mismatch increases the likelihood of severe complications for patients [4, 5].

The role of *killer-cell immunoglobulin-like receptor* (*KIR*) genes on *HSCT* outcome is not yet well understood, although several studies report an influence of donor *KIR* genotype on long-term survival after transplantation [6, 7]. With 17 genes, extensive gene copy number variation and 1,110 (Dec. 2020) described alleles in the *IPD-KIR* database, the *KIR* region also harbours formidable genetic diversity.

Large-scale sequence-based *HLA* and *KIR* genotyping is performed routinely for stem-cell donor registries and in clinical laboratories. Due to the diversity and complexity of these regions, high-resolution genotyping of the relevant genes crucially depends on the quality and comprehensiveness of the reference sequence databases [8, 9].

At present, for only 30% of the known *HLA* alleles the full genomic sequence is known, and the reliable reference-grade characterisation of genomic sequences of newly discovered *HLA* and *KIR* alleles remains technically challenging. The non-coding regions of these genes may contain extensive homopolymer tracts, i.e., stretches of single nucleotide repeats, or short tandem repeats [10]. Some genes may harbour structural indel variation, leading to differences of up to several kb in length between two alleles in a single heterozygous individual (e.g., Intron 1 of *HLA-DRB1*03:01:01:01*, 7,994 bp, and Intron 1 of *HLA-DRB1*07:01:01:01*, 10,281 bp, differ by 2,287 bp). Especially the *KIR* region contains large repeats, inversions and low-complexity regions. Additionally, due to extensive gene copy number variation in *KIR*, a single individual may accommodate three or more alleles for specific *KIR* genes.

Here, we present `DR2S` (**D**ual **R**edundant **R**eference **S**equencing), a tool designed to facilitate generating full-length phase-defined haplotype sequences in reference quality. While `DR2S` has been tested extensively on and optimised for *HLA* and *KIR* genes, it can be applied to any locus. Our approach takes advantage of the respective strengths of two readily available types of sequencing platforms: the accuracy of Illumina short-read sequencing and the read lengths achievable by third-generation single-molecule sequencing platforms.

While short-read sequencers typically produce highly accurate sequences, the length of a read is limited to about 300 bp. This is in many cases not sufficient to correctly phase allelic variants and thus results in ambiguous genotypes. Third-generation single-molecule sequencing technologies such as nanopore sequencing by ONT (Oxford Nanopore Technologies, Oxford, United Kingdom) or SMRT sequencing by PacBio (Pacific Biosciences, Menlo Park, California) are able to produce contiguous reads of several thousand base pairs. Yet, sequencing accuracy on these platforms is still severely limited and per-read error rates of up to 10% to 15% are common, especially in regions rich in homopolymers and repeats.

Currently, `DR2S` utilises data from targeted experiments, i.e., sequencing of full-length amplicons of the genes of interest. Separate fastq files from each sequencing experiment in combination with a generic reference sequence serve as input for `DR2S`.

### Implementation

`DR2S` is implemented as an R package [11]. It relies heavily on `Bioconductor` [12] and requires `bwa` [13], `minimap2` [14] and `IGV` [15] to be installed on the system. The use of system-wide installed `samtools` [16] is recommended, but the `Rsamtools` package may be used as a fallback [17]. `DR2S` is open source and available from GitHub (https://github.com/DKMS-LSL/dr2s). The framework of the `DR2S` pipeline and its major modules are described below. All mappings of short-reads are carried out using the `mem` algorithm of `bwa` whilst long-reads are mapped using `minimap2`.

#### Setup

The starting points for a `DR2S` analysis are gene-specific long-reads (PacBio or ONT) and short-reads (Illumina) of one or more samples provided as fastq files and a *generic reference sequence* of the gene that is analysed. It is also possible to rely exclusively on long-reads for haplotype separation and consensus calling, although this might not allow resolving repeat regions or homopolymers at a quality sufficient for submission to a reference database. In the case of *HLA* and *KIR* genes, providing the locus name as part of the initial run configuration is sufficient, for other genes a fasta file containing a reference sequence is required. All steps of the analysis workflow can be configured interactively in R or via YAML or JSON configuration files.

#### Filtering and Variant Definition

In a first step, a *sample-specific reference sequence* is created by mapping the short-reads to the generic reference sequence and calling the consensus (Fig. 1A). In this step, it is possible to reduce the sequencing coverage by sub-sampling the reads. The sub-sampling step is applied after the initial mapping and reads are sampled based on the coverage, and not on the number of reads alone. Consensus sequences are always inferred from the mapping by extracting the consensus matrix and subsequently calling a majority-rule-based consensus sequence.

**Figure 1.**
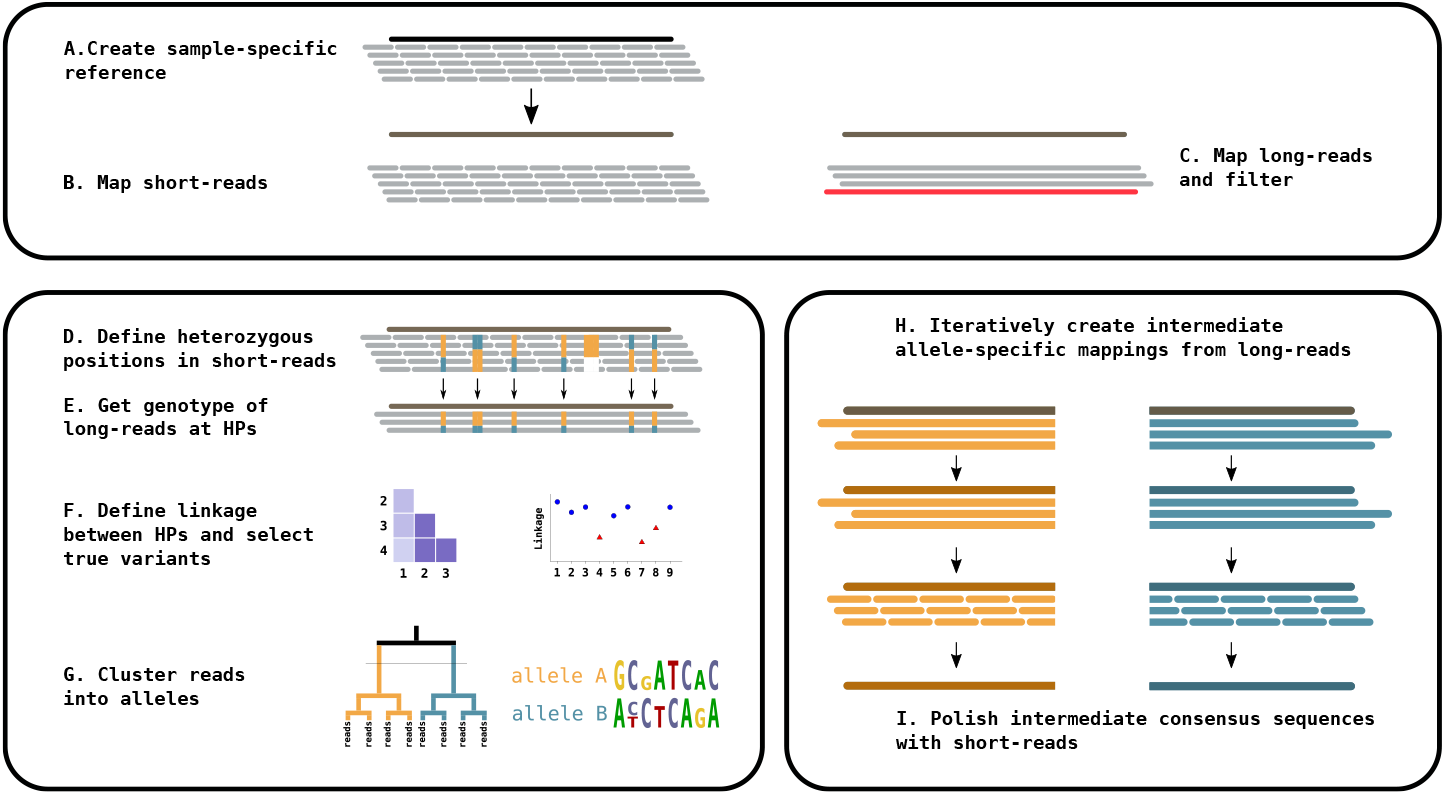
Workflow. Top panel: **(A)** The short-reads are mapped to the generic reference and a consensus is called to serve as a sample-specific reference. Short-reads **(B)** and long-reads **(C)** are mapped to the sample-specific reference. *Left panel*: Heterozygous positions (HPs) are defined in the short-reads **(D)** and used to infer the genotype at these positions in the long-reads **(E)**. ”True” HPs are distinguished from artefacts by linkage analysis **(F)** and only true HPs are clustered into alleles **(G)**. *Right panel*: Haplotype-specific long-reads are used to iteratively define draft haplotype consensus sequences **(H)** which are polished by the short-reads to call final haplotype consensus sequences **(I)**.

Next, both, long-reads and short-reads, are re-mapped to the sample-specific reference sequence (Fig. 1B and C, Fig. 2).

**Figure 2.**
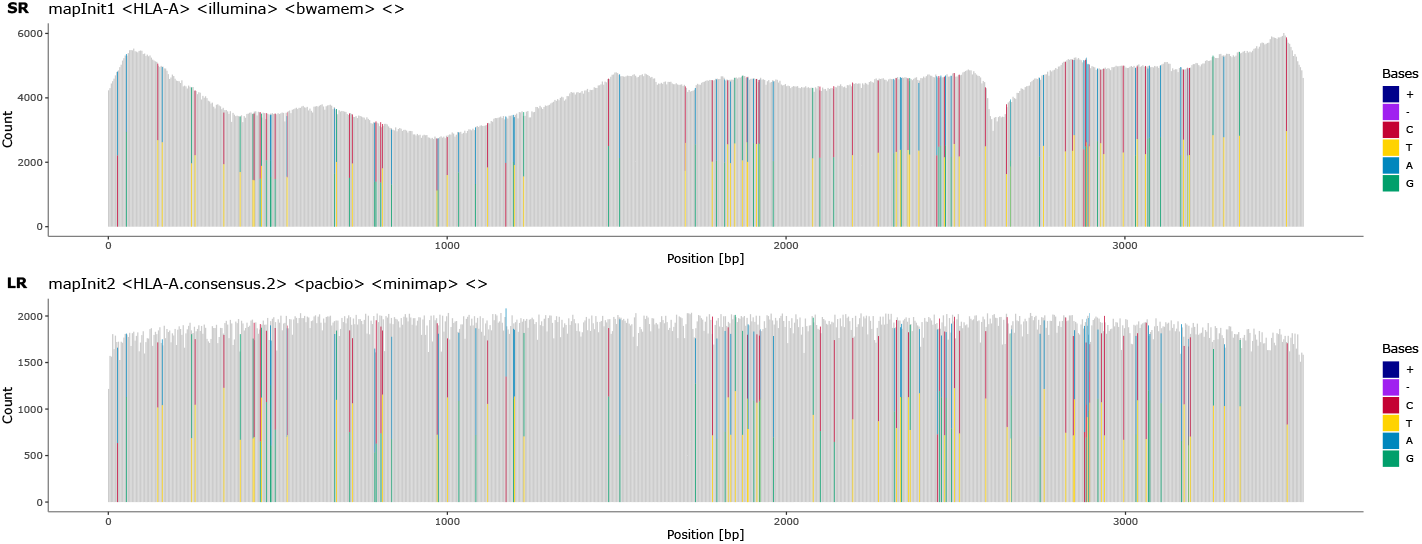
Read coverage and heterozygous positions in the primary mapping. The initial coverage of short-reads (**SR**) and long-reads (**LR**) against a generic locus-specific reference. Heterozygous positions (HP) and indels are colour-coded. HPs are shown by two colours, where both should, ideally, cover half of the height at a position. In this example, most HPs are present at the same position in both short-reads and long-reads. Observations of HPs that differ between short- and long-reads are common and are usually caused by differing allele imbalances between the two sequencing experiments. The clustering into two haplotypes is based on these HPs.

Optionally, the long-read mapping may be used to winnow out low-quality long-reads, where ”read quality” is assessed as similarity to a Position Weight Matrix (PWM) derived from the same mapping. The short-read mapping is used to infer the coordinates of non-gap heterozygous positions (HPs) with a minor allele frequency above a defined threshold (default 0.2; Fig. 1D).

The short-read-derived HP coordinates are then used to pinpoint heterozygous positions in the long-read mapping (Fig. 1E). The genotype at each non-gap HP is inferred for each long-read separately. Long-reads which do not cover at least 90% of HPs and HPs which are not covered by at least 30% of the long-reads are discarded.

Some HPs may be sequencing or mapping artefacts and should thus not be used for allocating reads to alleles. Such non-informative HPs are identified by linkage analysis (Fig. 1F), where linkage is measured as Cram’r’s V between all pairs of HPs. The matrix of pairwise linkage measures is used to cluster HPs into two groups, one with high intra-cluster linkage and the other with low intra-cluster linkage (Fig. 1F left and Fig. 3). HPs in the group with lower linkage are excluded if the mean intra-cluster linkage values between the two groups differ by more than a set threshold (Fig. 1F right and Fig. 4).

**Figure 3.**
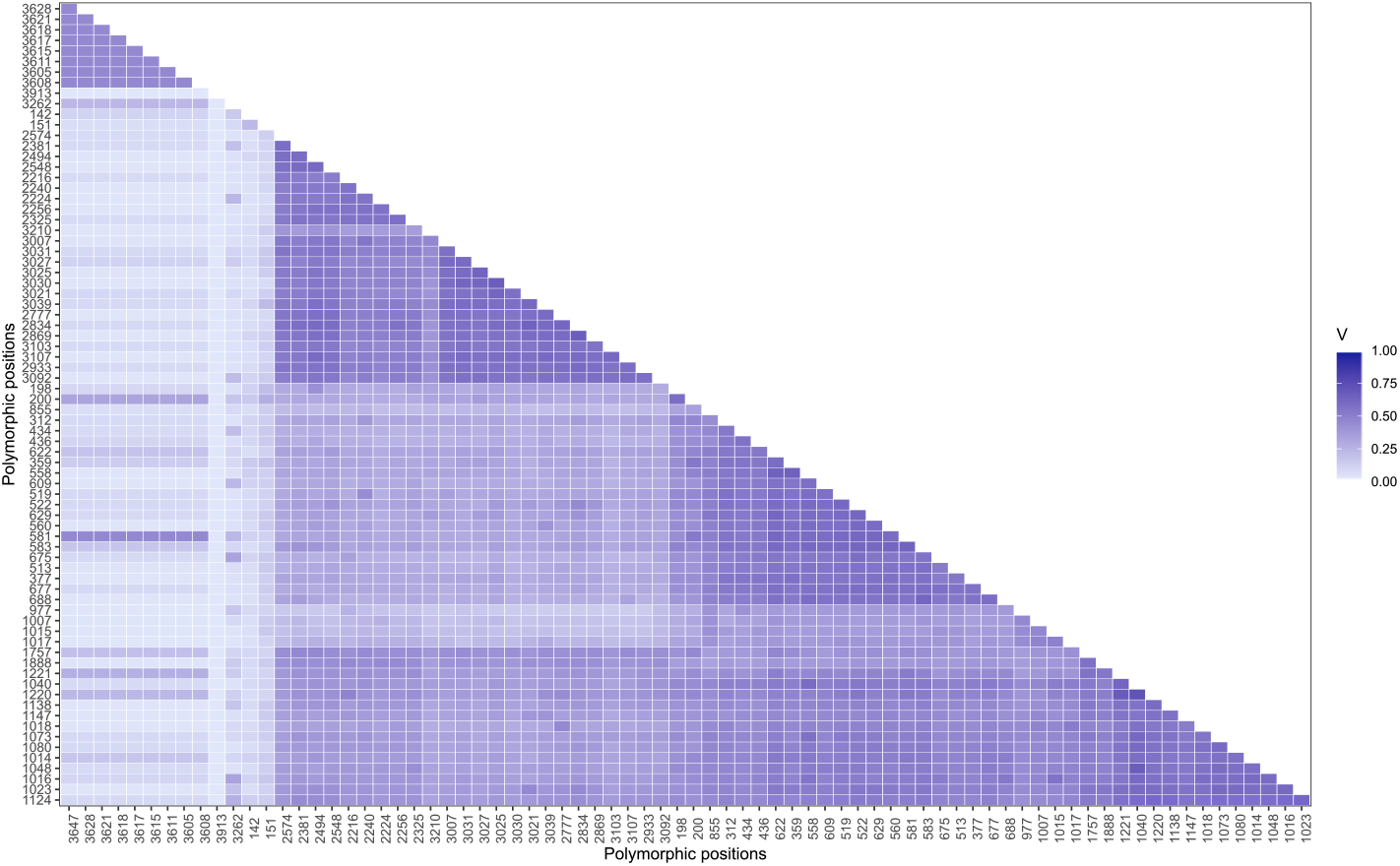
Linkage of heterozygous positions. Correlation matrix of pairwise linkage between all HPs. The order of positions in this plot is based on a hierarchical clustering of pair-wise distances. Positions with a low correlation to many other positions (indicated by a light color) are most likely sequencing artefacts and excluded from long-read clustering.

**Figure 4.**
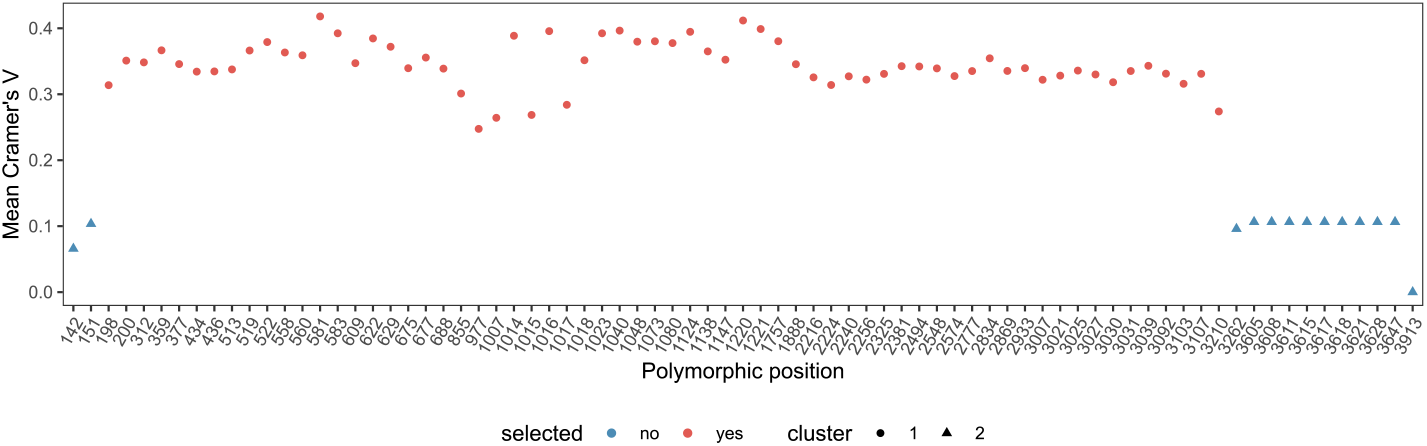
Linkage of heterozygous positions. Linkage between HPs measured by Cramér’s V. Several potential HPs are not linked to other HPs and are thus excluded from long-read clustering.

Based on the remaining long-reads and HPs, reads are categorised into *haplotype-specific long-read sets* (Fig. 1G). The number of possible haplotypes is not limited to two, as expected for single-copy heterozygous genes, but DR2S can also deal with cases of multiple gene copies as encountered, for example, in the *KIR* region or in polyploid organisms.

#### Long-Read Clustering

To generate haplotype-specific long-read sets, the genotype of each HP in each long-read is used to construct a Position-Specific Distance Matrix (PSDM). A PSDM can be derived from a PWM by weighting the distance between two reads by the nucleotide weights at each polymorphic position such that differences in major genotypes at a position count more towards distance than differences between major and minor genotypes. As an example, consider a heterozygous position with a distribution of 50% A, 45% G and 5% T nucleotides. Here, the position should contribute more heavily to the overall sequence distance between two reads if the reads feature A and G, respectively, while a T is more likely to derive from a sequencing error.

Hierarchical clustering is applied to the PSDM and the cut height, i.e., the most likely number of clusters is inferred using adaptive branch pruning as implemented in the *dynamicTreeCut* package [18]. If this approach yields more clusters than the expected number of alleles, only the most distant clusters are retained. All reads are then re-scored with respect to the PWMs derived from the retained clusters generating *haplotype membership coefficients*. Finally, only a fraction of reads best representing each cluster based on *haplotype membership coefficients* are retained for further processing (see Fig. 5). This strategy effectively eliminates chimeric reads formed during PCR and other amplification or sequencing artefacts from interfering with downstream haplotype reconstruction, as long as chimeric reads stay less abundant than reads true to the actual alleles present in a sample.

**Figure 5.**
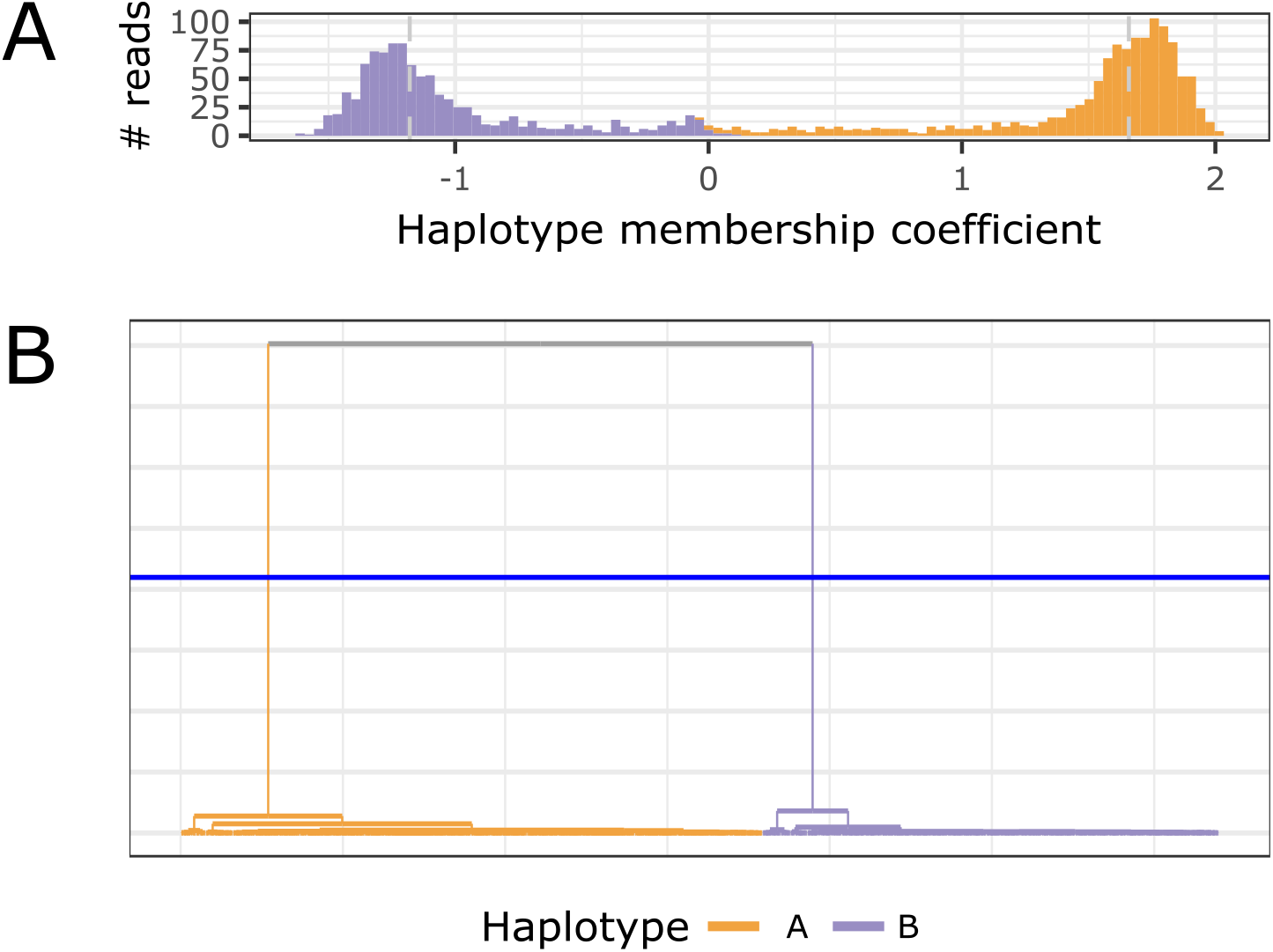
Long-read clustering. **A** Histogram of the *haplotype membership coefficient* of clustered long-reads. A negative value suggests a read membership in haplotype B, a positive value suggests a read membership in haplotype A. The dashed vertical lines mark heuristically determined thresholds for retaining long-reads to iteratively generate haplotype-specific consensus sequences. **B** Hierarchical clustering dendrogram. Each leaf represents a long-read. The blue horizontal line indicates the adaptively chosen cut-height for separating reads into two clusters.

#### Consensus Calling

The long-reads retained for each cluster are now considered to be derived from distinct alleles and stored in separate fastq files to serve as input for generating draft *haplotype-specific reference sequences*.

For that purpose, each haplotype-specific long-read set is mapped iteratively to a consensus sequence created at the previous iteration (Fig. 1H). The consensus sequence for the first iteration is derived from the initial mapping step by extracting a consensus matrix from the mapping and keeping only reads of the haplotype. The iterative refinement of the consensus sequences allows the resolution of haplotype-specific indel variants. Two iterations are generally sufficient for long-reads to converge on a draft reference sequence (see Fig. 6). In the final step, these draft references are corroborated or polished as necessary using the short-read data.

**Figure 6.**
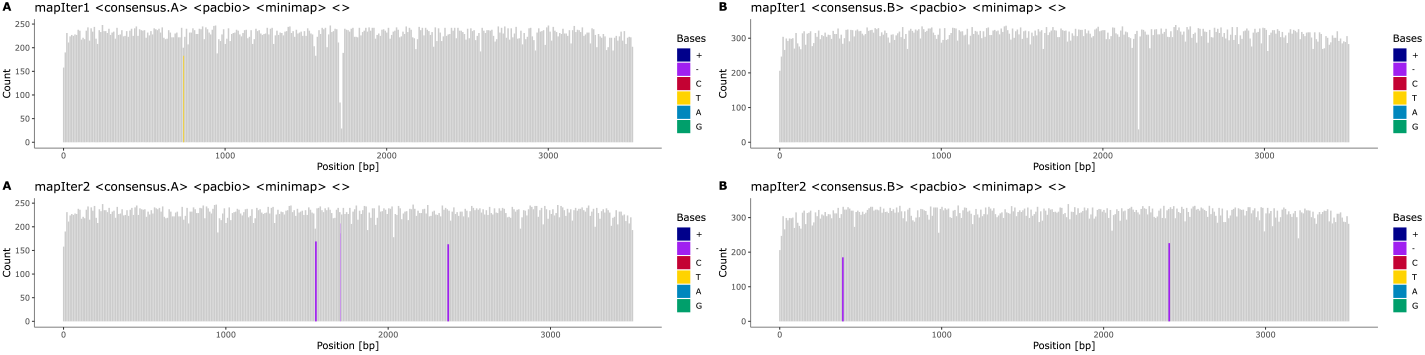
Iterative mapping of long-reads. Haplotype-specific references for alleles **A** and **B** are created iteratively. While some ambiguities remain in the long-read pileup after an initial round of mapping (**mapIter1**), alignment accuracy is significantly improved by remapping the reads sets against consensus sequences derived from the first mapping (**mapIter2**). The majority of HPs in the mapping are resolved at the end of the iterative mapping (lower panel). Some positions still harbour deletions, denoted by the purple bars, which need to be resolved by short-reads in the final mapping step.

#### Consensus Polishing

To polish the long-read-derived haplotype-specific draft reference sequences, short-reads are also classified based on their putative haplotype of origin.

To that end, short-reads that cover an HP are assigned to a haplotype cluster based on the respective long-read cluster. Short-reads that do not cover HPs are distributed to all haplotype clusters. Finally, short-reads are mapped to the long-read-derived haplotype-specific draft reference sequences obtained in the previous step. The final consensus sequences are derived from these short-read mappings (Figs. 1I and Fig. 7). Remaining ambiguities that are covered by clustered short-reads reads should be resolved after this step.

**Figure 7.**
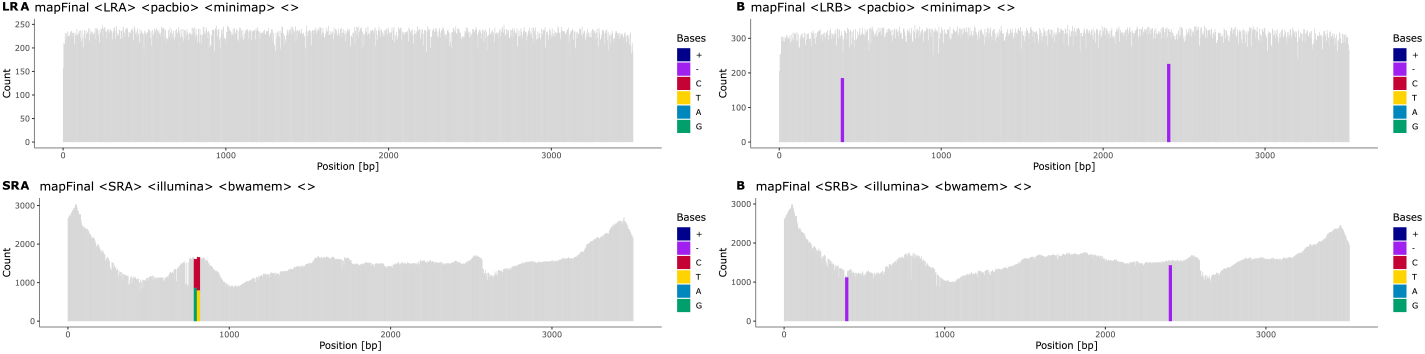
Final mapping of long-reads and short-reads to the refined, haplotype-specific references. Heterozygous positions (HP) and indels are colour-coded. Purple bars in the coverage plot (e.g. at position ∼ 400 bp in haplotype B) visualise deletions that are present in the short-read mappings of both haplotypes and indicate spurious insertions or positions that are hard to resolve using long-reads alone. Bicoloured positions denote existing ambiguities in the final mapping, in this example caused by positions where short-reads are not clustered with the two haplotypes as expected (e.g. at position 790 in haplotype A). It is still possible to resolve such positions using long-reads, even if they were not used for the initial haplotype clustering. The final consensus sequences are derived from these mappings.

#### Reporting and Editing

During a run, statistics and diagnostics plots are created at each step to aid evaluating the quality of haplotype reconstruction. A coverage plot is created for each mapping, highlighting heterozygous positions, indels and positions that do not match the reference (Fig. 2, Fig. 6 and Fig. 7). Each step of the long-read clustering is documented by plots such as the dendrogram of the hierarchical clustering analysis and the sequence logo of the HP matrix for each haplotype (Fig. 8). Potential problems and artefacts, such as positions that remain heterozygous in the final haplotype-specific mappings or longer insertions that cannot be resolved automatically, are reported and may need to be corrected manually. To aid manual correction, configuration files for IGV are created that allow quickly visualising the long-read and short-read mappings at positions deemed ambiguous by the software (Fig. 9). The consistency of homopolymer runs is checked separately for positions that exceed a configurable homopolymer run length. Here, distributions of homopolymer lengths over individual reads are calculated and plotted for each haplotype, and the mode value is compared to the homopolymer length of the final consensus sequence of the respective haplotypes (see Fig. 10).

**Figure 8.**
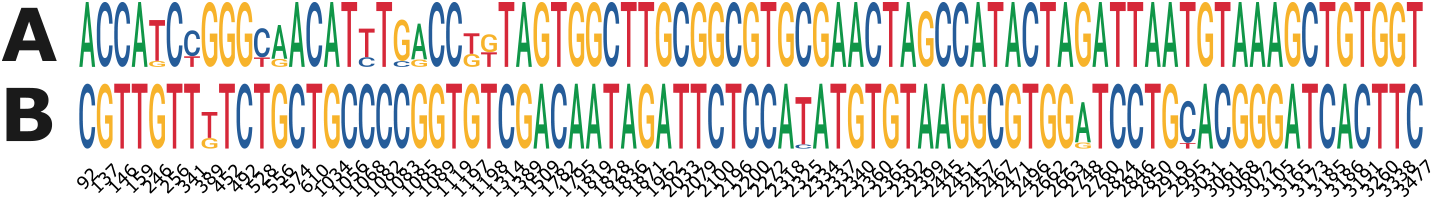
Sequence logo of heterozygous positions. Most positions clearly distinguish the two haplotypes **A** and **B**. Only few positions in both haplotypes are of low bit-wise information content.

**Figure 9.**
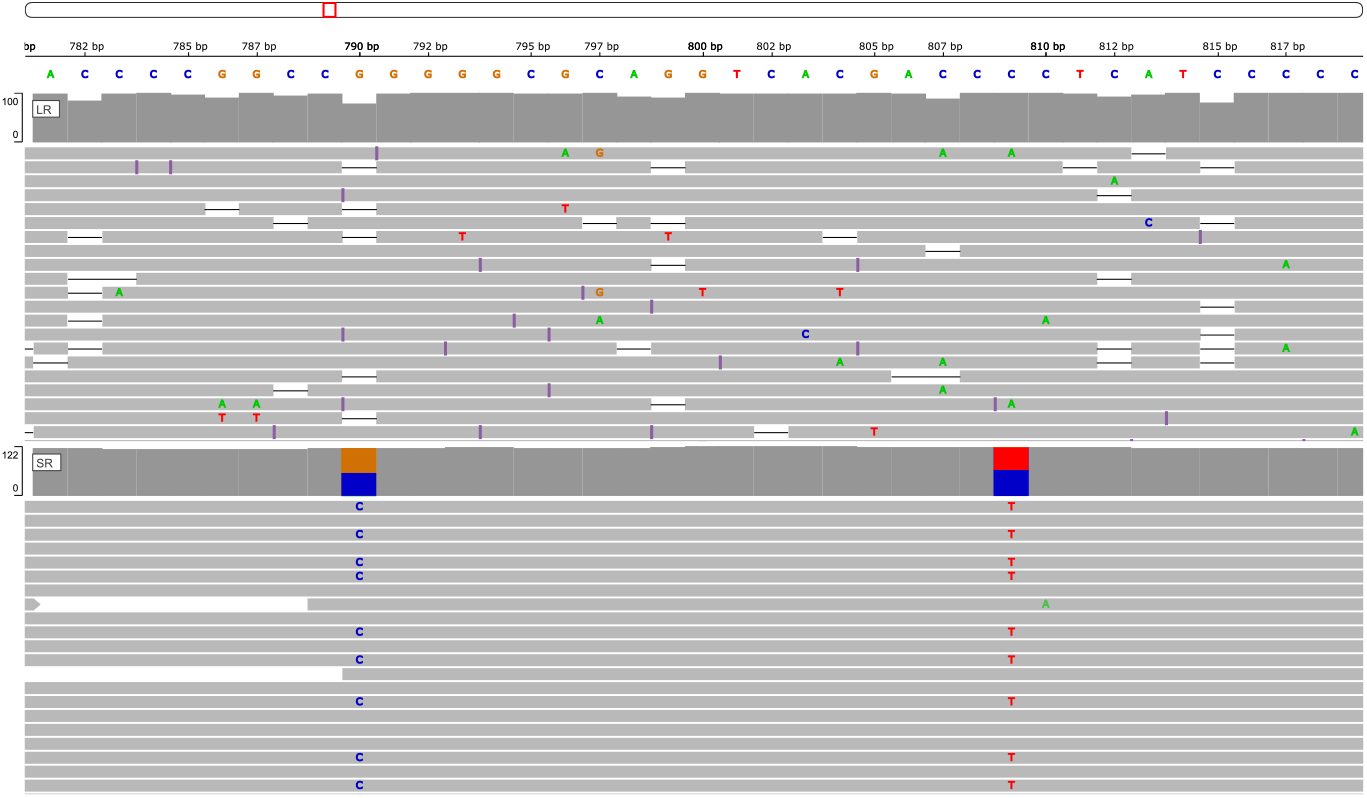
Alignments of long-reads and short-reads against one inferred haplotype. This screenshot from `IGV` (Integrative Genomics Viewer) shows PacBio reads (upper panel) and Illumina reads (lower panel) aligned to one of the inferred haplotype sequences prior to manual correction. The alignments are easily visualised using configuration files automatically generated by `DR2S`. This example demonstrates remaining ambiguities in the short-reads at positions 790 and 809. These positions were also observed in the final mapping plot. The longreads at these positions are unambiguous and can be used to guide the manual edit.

**Figure 10.**
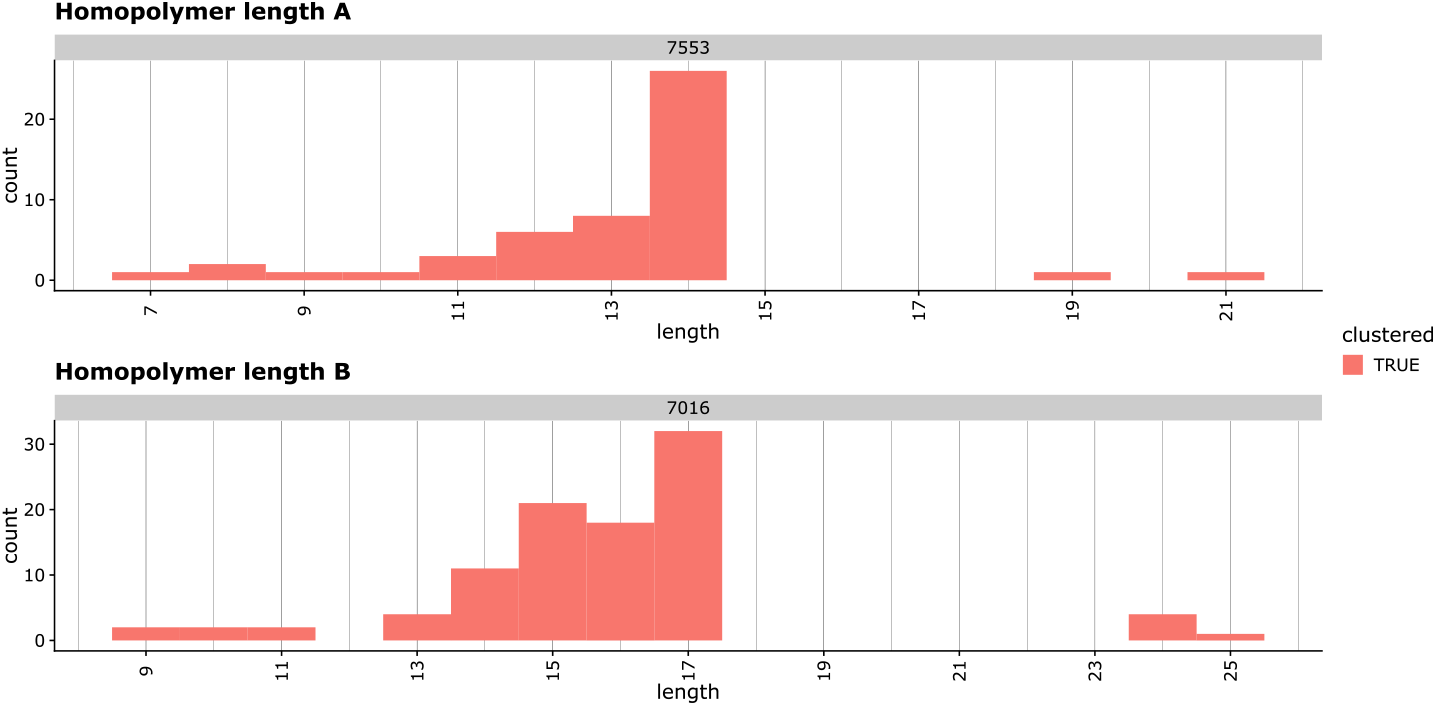
Example of homopolymer visualisation. Short-reads that cover a homopolymer exceeding a set length are visualised by histograms. The true length of a homopolymer is usually the mode value, i.e. the length supported by most reads. In this example, haplotype A has a homopolymer length of 14 at position 7553, while haplotype B has a homopolymer length of 17 at position 7016.

Manual edits are made in a preliminary alignment file of the haplotype consensus sequences (Fig. 11). To evaluate the effects of manual edits, functions are provided to remap haplotype-specific reads to the updated references (Fig. 12) and to visualise the remapping results. This allows for a straightforward iteration over problematic positions that the software could not resolve automatically. Once the user has asserted the correctness of the haplotype reference sequences, they can be ”checked out” into final *fasta* files.

**Figure 11.**
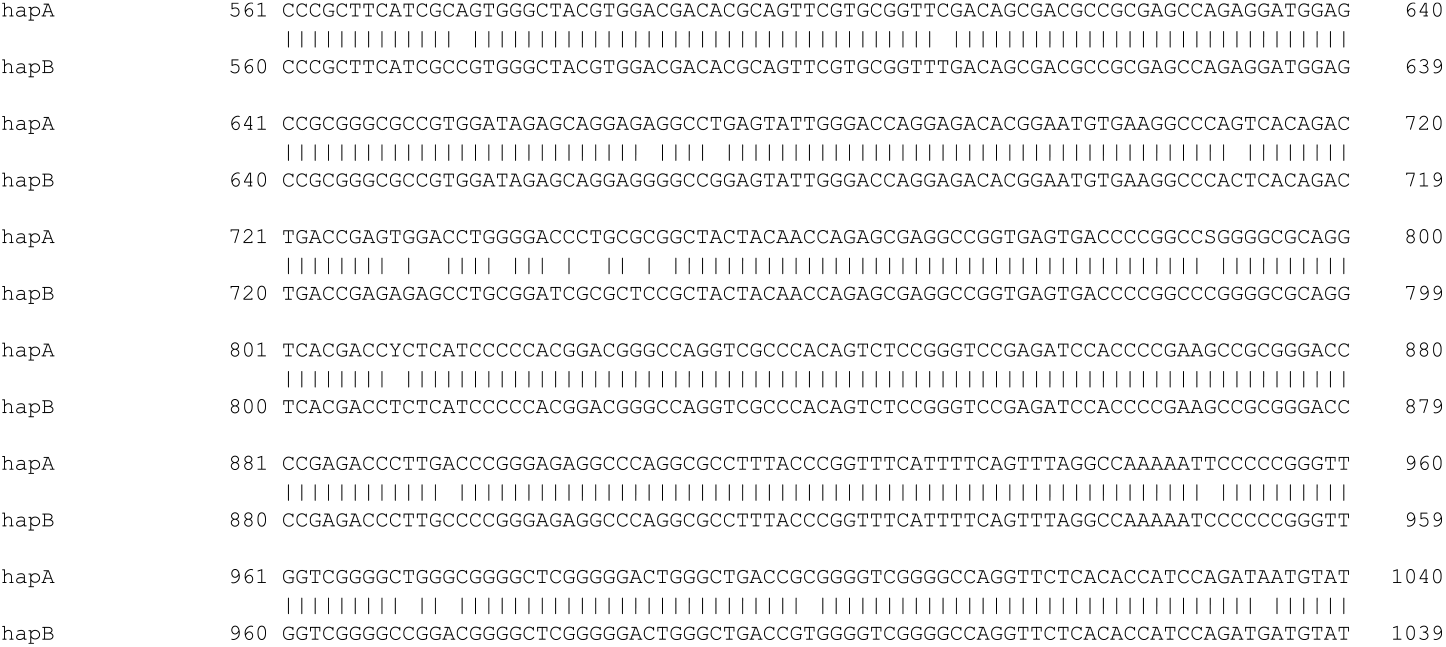
The preliminary alignment of all haplotypes. Sequences of all haplotypes can manually be edited in a text editor and can be checked by remapping reads to the changed sequence. In this example, ambiguous positions at positions 790 and 809 in haplotype A are denoted by the IUPAC codes S and Y, and need to be manually checked and corrected.

**Figure 12.**
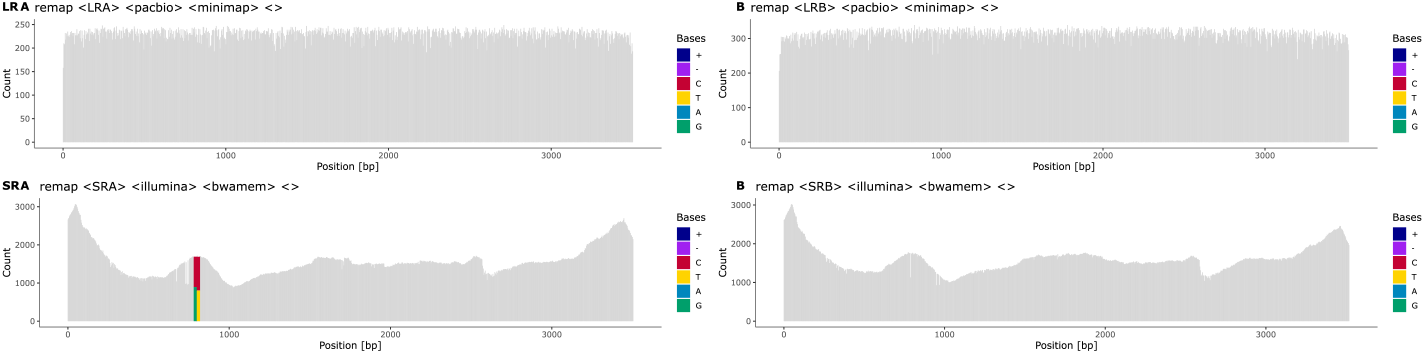
The remapping of short-reads and long-reads to manually updated references. All heterozygous positions and indels have been resolved and the final sequences are fully phased. The ambiguous positions in haplotype A are still shown as the composition of reads is not changed during the reporting, but the long-reads show a correct consensus sequence.

## Results and Discussion

We developed `DR2S` specifically to address the challenge of assembling and ascertaining novel *HLA* and *KIR* allele consensus sequences using easy-to-generate next-generation sequence data from heterozygous samples. We have been using DR2S routinely and successfully to create several hundred high-quality, fully-phased, reference allele sequences for *HLA* and *KIR* genes for submission to the *IPD-IMGT/HLA* and *IPD-KIR* databases [19].

We compared the performance of `DR2S` to two existing haplotype assembly tools, `WhatsHap` [20] and `HapCUT2` [21]. These tools were chosen based on their ability to utilise both short-read and long-read genomic data as input and their reported performance relative to alternative solutions. Note that both `WhatsHap` and `HapCUT2` require BAM/CRAM files containing reads aligned to a genomic reference and a VCF file containing corresponding variant calls (SNVs and indels) as input. It is left to the user to generate these input data from unmapped sequence data, although a recommended workflow exists for `WhatsHap` (https://whatshap.readthedocs.io/en/latest/guide.html).

We evaluated the three phasing tools using *HLA* sequence data created by long-range whole-gene amplification followed by fragmentation for shotgun sequencing on an Illumina MiSeq instrument and direct long-read sequencing on PacBio’s Sequel II and ONT’s MinION platforms with R10.3 flow cells, respectively, as described previously [19]. Five samples were sequenced for six *HLA* genes (*HLA-A, -B, -C, -DRB1, -DQB1*, and *-DPB1*) on the Illumina and ONT platforms. Five additional samples were sequenced for five *HLA* genes (*HLA-A, -B, -C, -DQB1*, and *-DPB1*) on the Illumina and PacBio platforms. To phase these samples with `WhatsHap` and `HapCUT2`, we followed `WhatsHap`’s recommended workflow, using `bwa mem` and `minimap2` for the initial short-read and long-read mapping, respectively, followed by variant calling on short-reads alone using the `FreeBayes` variant caller [22]. All analyses were carried out using the default parametrisation of the respective tool.

Since for none of these samples independent allele sequence data were available, we established ”ground truth” haplotype sequences by performing an initial run of `DR2S` followed by a careful visual inspection and manual curation of all resulting allele sequences. We used these curated sequences as the basis for calculating the error rates of each tool without manual curation. The haplotypes assembled by each tool for each sample and gene were attributed to their target ground truth haplotype by overall similarity. The accuracy of the haplotype reconstruction was assessed using *mismatch error rate* and *phase switch error rate* as metrics. We defined *mismatch errors* as single variants or indels not matching the target ground truth haplotype nor attributable to the alternative ground truth haplotype, and *phase switch errors* as single variants or runs of consecutive variants attributable to the alternative ground truth haplotype per sample per gene. Error rates were calculated by considering the total number of deviating positions of a ground truth haplotype to the gene-specific generic reference sequence used in the initial mapping step of each tool, as the maximum number of possible errors. For both, PacBio and ONT long-reads, DR2S reconstructed the most accurate haplotypes both with respect to mismatch and phase switch errors (Fig. 13).

**Figure 13.**
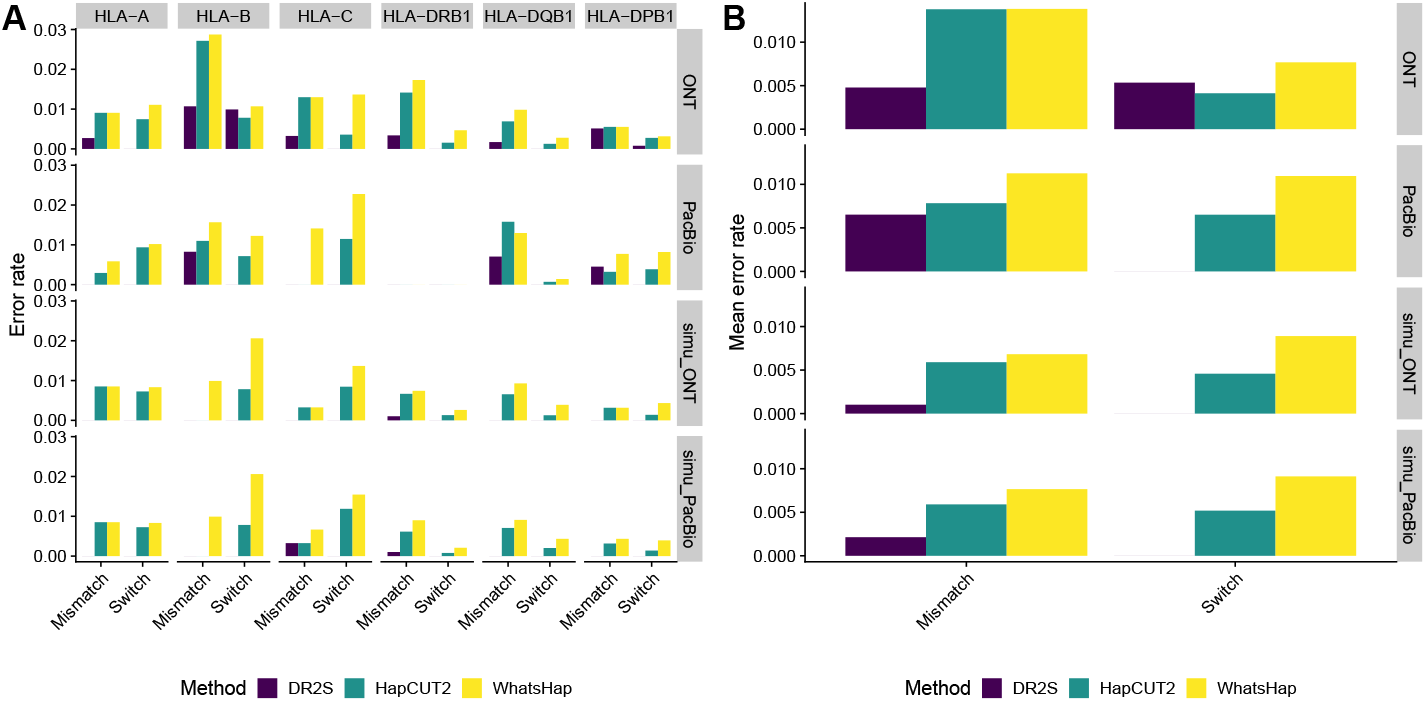
Benchmarking DR2S against two other haplotype assembly methods. Mismatch and phase switch error rates for different *HLA* genes (A) and average error rates across all genes **(B)**. The panels show results for ONT long-reads in combination with Illumina short-reads (ONT), PacBio long-reads in combination with Illumina short-reads (PacBio) and simulated ONT or PacBio long-reads in combination with simulated Illumina short-reads (simu ONT and simu PacBio). Note that different samples were used with both long-read sequencing technologies and that no PacBio long-read data were available for *HLA-DRB1*.

In addition to real HLA sequence data sets, we also compared the performance of `DR2S`, `WhatsHap` and `HapCUT2` using simulated sequencing data. This should mitigate the potential biases arising from using `DR2S` for creating ground truth sequences in the first place. To simulate benchmark data we used the ground truth sequences derived from the real data sets as seeds. Simulations of Illumina MiSeq data were carried out using `InSilicoSeq` [23] with a targeted read depth of 2000 and the *miseq* model file provided. PacBio and ONT data were simulated with `PBSIM2` [24]. We used the *P5C3* model for PacBio data and *R10*.*3* for ONT data, respectively. The parameters for sequencing error ratios were used as suggested by `PBSIM2`, i.e. a mean accuracy of 85% and a substitution/insertion/deletion ratio of 6/50/54 and 23/31/46 for PacBio and ONT, respectively. Both tools were chosen for their ability to simulate readswithout the need to first build empirical error models based on supplemented sequence data, which again might have introduced a bias towards DR2S. Again, for both, simulated PacBio and simulated ONT long-reads, `DR2S` delivered the most accurate haplotypes of all three tools with regard to mismatch and phase switch errors (Fig. 13). For DR2S, we found that all remaining mismatching positions were flagged by the tool as potentially problematic, thus facilitating manual curation.

All tools exhibited comparatively large heterogeneity in error rates across *HLA* genes, especially with the real data sets (Fig. 13A). This likely reflects the fact that data quality across samples and loci varied widely, especially with regard to read coverage (compare Fig. 14), the sequence complexity of the specific alleles found in a sample, and the distance of specific alleles to the reference sequence used for the initial mappings. Overall, we found little difference in error rates with regard to the two long-read sequencing technologies used (Fig. 13B). This indicates that the generally larger per-read error rates of nanopore reads relative to PacBio reads are not necessarily an impediment to accurate haplotype reconstruction, at least if used in conjunction with highly accurate short-reads.

**Figure 14.**
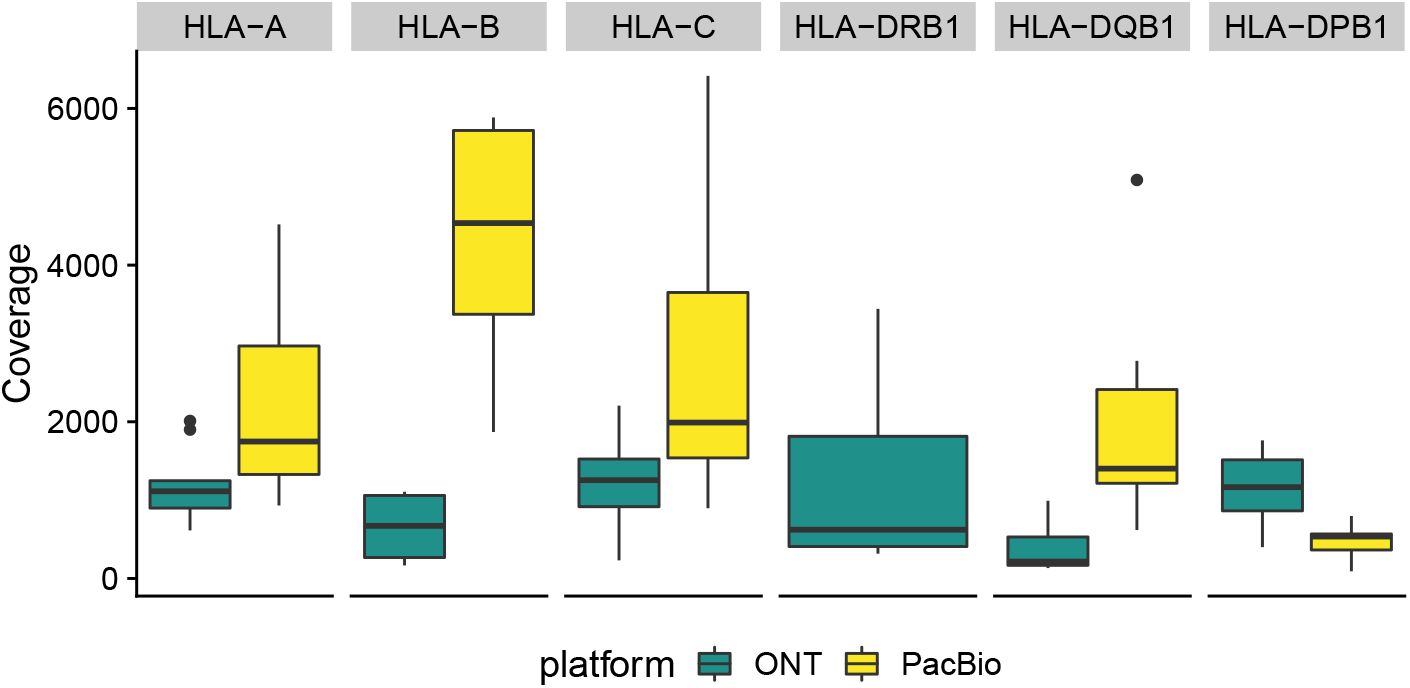
Coverage of the benchmarked samples. Boxplot of observed sequencing coverage by gene and sequencing platform.

Overall, even without manual curation, `DR2S` showed a reduction of approximately 60% in mismatch error rate over both `HapCUT2` and `WhatsHap` (Fig. 13B). This is likely due to the heavy emphasis put by `DR2S`’s workflow on iteratively refining the reference used for read mapping, and the aggressive pre-selection of sequence reads used for haplotype reconstruction according to their *haplotype membership coefficient* (compare Fig. 5). Thereby `DR2S` arguably achieves at a cleaner base for assembling the two haplotypes harboured by a heterozygous sample. In contrast, `HapCUT2` and `WhatsHap` leave it to the user to provide a read mapping and a set of variants to be phased. The recommended workflow for these tools does not provide any guidelines as to how to prepare or pre-process the sequence data for potentially improved results.

Moreover, in contrast to `HapCUT2` and `WhatsHap`, `DR2S` exhibited no phase switch errors for any of the analysed samples (Fig. 13). Again, this is likely due to the read pruning strategy employed by `DR2S` when assigning long-reads to allele clusters, which effectively eliminates or reduces artefacts such as PCR chimeras or low-quality reads that otherwise may interfere with haplotype assembly.

These results demonstrate that `DR2S` can be used to create haplotype sequences for highly polymorphic genes such as the *HLA* genes with very high accuracy. However, it is also clear that for any haplotype assembly tool, depending on the raw data quality and the complexity of the region of interest, occasional errors will be introduced. If the aim is to submit the resulting consensus sequences to a database, confidence in the resulting sequences is of particular importance and some degree of visual sequence validation is inevitable. Extant tools do not explicitly cater for this need, thus requiring expert bioinformatics knowledge to create custom workflows for inspection and validation of their results. In contrast, `DR2S` implements a number of post-processing features to alert the user to potentially dubious positions, to facilitate visual inspection of the alignment data using preconfigured `IGV` plots, and to iteratively edit and re-evaluate haplotype reference sequences. The easy manual inspection of mappings and problematic positions ensures highly trustworthy final sequences.

In our environment, `DR2S` is used in a compute cluster and a single sample/gene usually needs between 5 and 20 minutes to finish on 8 cores, depending on sequencing coverage and gene length.

## Conclusions

`DR2S` is a largely automated workflow designed to create high-quality fully-phased reference allele sequences for highly polymorphic gene regions such as *HLA* or *KIR*. Designed to work with a combination of short-read and long-read amplicon data from a region of interest, it shows superior performance to comparable tools both in terms of mismatch errors and phase switch errors. In addition, `DR2S` offers supporting tools to appraise the quality of the resulting haplotypes, perform manual edits, and assess the consequences of these edits. `DR2S` has been used by biologists to successfully characterise and submit more than 500 *HLA* alleles and more than 500 *KIR* alleles to the *IPD-IMGT/HLA* and *IPD-KIR* databases.

## Availability and requirements

### Project name

DR2S

### Project home page

https://github.com/DKMS-LSL/dr2s

### Operating system(s)

Linux

### Programming language

GNU R

### Other requirements

Samtools, BWA, minimap2, IGV

### License

MIT

### Any restrictions to use by non-academics

None

## Abbreviations

HLA: Human Leukocyte Antigen
KIR: Killer-cell Immunoglobulin-like Receptor
HSCT: Haematopoetic stem-cell transplantation
HP: Heterozygous Position

